# Natal philopatry, dispersal and age of first breeding in relation to size and sex of Arctic Terns *Sterna paradisaea*

**DOI:** 10.1101/2025.05.22.655618

**Authors:** Chris P.F. Redfern, David Steel, Paul G. Morrison

## Abstract

Many seabird species are in decline and population demographic models are important for revealing the causes and developing conservation strategies. Natal and breeding dispersal are key parameters of such models but can be challenging to estimate and may vary by sex. Along the Northumberland coast, Arctic Terns Sterna paradisaea nest across sites separated by distances up to 32 km. From ringing and recapture of nestling and nesting adult Arctic Terns over two decades, natal philopatry in component sites of this metapopulation was high and recruitment to a colony with managed public access was similar to nearby colonies with no public access. Mean head length of nesting birds recruited from non-natal sites was significantly smaller than those nesting on their natal site. Sexual-size dimorphism was used to estimate the proportions of each sex in capture samples and indicated that males were generally faithful to their natal site but up to nearly a third of females may have dispersed to non-natal sites. Arctic Terns breed from two years of age; head length data indicated that breeding birds of two to four years old were mainly female, and suggested that the first-breeding age of males was up to three years older. Young breeding birds were caught later in breeding seasons than older birds. Unexpected colony abandonment can confound estimates of natal philopatry and dispersal in metapopulations. These results demonstrate the value of mark-recapture studies and indicate that sex-specific dispersal and breeding-age parameters will be essential components of demographic models.

---

Seabirds are uniquely adapted to the oceanic environment covering 72% of the earth’s surface but are in decline worldwide as a consequence of climate change, overfishing and fisheries bycatch (Dias et al. 2019, Melvin et al. 2023, Ramírez et al. 2024). Monitoring breeding numbers and productivity annually at seabird colonies is essential for identifying trends and developing demographic models for understanding the reasons for change and designing conservation strategies. From genetic, morphological and speciation perspectives, seabird population structures vary from panmixia to fragmentation into distinct units with limited breeding ranges despite wide distributional ranges at oceanic scales (Friesen et al. 2007, Garrett et al. 2020). Demographic models integrating migratory and metapopulation dynamics can also help understand the evolutionary processes that have driven and may drive future speciation and subspeciation processes in seabirds. In this context, the demography of Arctic Terns *Sterna paradisaea* is of particular interest. This species has possibly the widest global range of any seabird, with a circumpolar breeding distribution in the Northern Hemisphere (Hatch et al. 2020), a nearly circumpolar non-breeding distribution around the Antarctic coastline (Salomonsen 1967) and, for Arctic Terns from northwest European populations at least, a migration from breeding to non-breeding areas that can involve a traverse across the entire southern Indian Ocean to East Antarctica (Redfern & Bevan 2022). In contrast, the closest relative of the Arctic Tern, the Antarctic Tern *Sterna vittata* (Bridge et al. 2005), is differentiated into distinct subspecies breeding on island sites distributed round the Southern Hemisphere (Connan et al. 2015).

Throughout their breeding range, Arctic Tern populations are in decline (Burnham et al. 2017, Mallory et al. 2018, Henri et al. 2020, Wyn 2023, BirdLife International 2024, Stanbury et al. 2024) with changes in food abundance (Suddaby & Ratcliffe 1997, Henri et al. 2020), predator pressure (Burnham et al. 2017, Scopel & Diamond 2017) and climate (Redfern & Bevan 2020, Morten et al. 2023) likely to be key factors. As an important component of seabird biodiversity and sensitive to the all-pervading impact of human populations, it is a species in need of conservation, especially at the limits of its range. To understand how changes in mortality and/or productivity translate to overall population growth, linkage between colonies which comprise part of a wider metapopulation must be incorporated into model structures to assess potential for adaptation to environmental challenges through dispersal and immigration (Gaya et al. 2024). Natal philopatry or dispersal is a critical component of demographic models (Border et al. 2017, Cantrell et al. 2017), facilitates gene flow between populations (Penttinen et al. 2024) and is important for management planning (Allen & Singh 2016).

Most Arctic Terns ringed in Britain and Ireland are ringed as nestlings (88.5%, 2002 to 2023) (Robinson et al. 2024). In Northumberland on the east coast of Great Britain, Arctic Terns are near the southern limit of their breeding range at these longitudes. Arctic Terns in this population have been ringed intensively for over 70 years, particularly on the Farne Islands where Arctic Terns have represented 26% of the ringing totals for this species across Britain and Ireland (author data in conjunction with Robinson et al. (2024)) and this has contributed to understanding of movements (Radford 1961). However, with a low recovery rate (Appleton et al. 1997), the ringing of nestling Arctic Terns has limited value for population studies without mark-recapture studies on breeding birds and for the last two decades nesting adults have been trapped on three island sites across Northumberland. The aim of this study was to investigate the connectivity of colonies within the Northumberland metapopulation because this is important for developing demographic models for conservation planning. Our objectives were to determine, from mark-recapture data, the extent of natal philopatry at each colony and whether this varied in relation to human activity, and to address the questions whether dispersal and age of first breeding varied in relation to sex.

## METHODS

### Study sites

In Northumberland, Arctic Terns nest predominantly on the Farne Islands (mainly Inner Farne, 2.5 km from the mainland, and Brownsman, 2.6 km northeast of Inner Farne), a mainland site, the Long Nanny, 9 km south of Inner Farne, and Coquet Island 32 km south of Inner Farne (Fig. 1a). Coordinates (decimal degrees; longitude, latitude) are: Inner Farne −1.6546, 55.6163; Brownsman −1.6256, 55.6341; Coquet Island −1.54°, 55.33°. Arctic Tern nestlings and adults have been ringed by the authors on Coquet Island from 1995, and the Farne Islands from 1996; others have ringed nestlings at the Long Nanny colony over the same period (Supplementary Information, Section 1). Adult Arctic Terns were trapped or retrapped (for techniques see Redfern & Steel (2024)) on the nest on Inner Farne (to 2019) and Brownsman (to 2014). Arctic Terns were also trapped on the nest on Coquet Island to 2015, and again from 2020 to 2022. The intensity of nest trapping varied by site and year and was done when opportunities allowed; the year ranges for captures of known-age adults at each site are summarised in Table 1.

**Table 1.** Known-age Arctic Terns trapped as nesting adults: colonies of origin and percentages from different Northumberland colonies of Inner Farne (INF), Brownsman (BRO), The Long Nanny (LNAN) and Coquet Island (COQ). Bold and underlined font indicates natal colony. Fisher’s Exact tests for proportions of natal (n) and non-natal (nn) Arctic Terns nesting at each site, using estimated totals corrected for expectations from the number of nestlings ringed at each site (see Methods): Inner Farne (n 157; nn 39) compared to Brownsman (n 20; nn 8), P = 0.32; Farnes (n 177; nn 47) compared to Coquet Island (n 60; nn 8), P = 0.11. Comparing Coquet Island with Coquet Island 2020 – 2022 (n 9; nn 15), P < 0.00001.

| Nesting adults (inclusive year range of captures) | Colony of origin |  |  |  |  |  | Percentage originating from Northumberland colonies<sup>¶</sup> |  |  |  |
| --- | --- | --- | --- | --- | --- | --- | --- | --- | --- | --- |
|  | INF | BRO | LNAN | COQ | other | Total | INF | BRO | LNAN | COQ |
| Inner Farne (2005-2019) | **<u>157</u>** | 12 | 3 | 8 | 3<sup>a</sup> | 183 | **<u>80.1</u>** | 10.7 | 2.6 | 6.6 |
| Brownsman (2007-2014) | 8 | **<u>20</u>** | 0 | 5 | 0 | 33 | 17.9 | **<u>71.4</u>** | 0 | 10.7 |
| Coquet (2000-2015)* | 5 | 1 | 3 | **<u>60</u>** | 1<sup>b</sup> | 70 | 5.9 | 1.5 | 4.4 | **<u>88.2</u>** |
| Coquet (2020-2022)* | 10 | 7 | 1 | **<u>9</u>** | 0 | 27 |  |  |  |  |
| Coquet (2020-2022)-adjusted<sup>¶¶</sup> | 6 | 8 | 1 | **<u>9</u>** | 0 | 24 |  |  |  |  |

<sup>a</sup>Isle of May (2); Port of Dublin (1); <sup>b</sup>One from Port of Dublin; <sup>¶</sup>corrected to the same number of pulli ringed in each of the originating colonies for the relevant year ranges from 1995 to: 2017 for Inner Farne captures, 2012 for Brownsman captures and 2013 captures with natal colony as baseline. The first possible year of recruitment will be in 3<sup>rd</sup> calendar year after ringing (two years old). <sup>¶¶</sup>Corrected for the number of pulli ringed in each originating colony up to and including 2019. \*Coquet: no nest trapping 2016-2019, but resumed in 2020-2022; none of the previous known-age birds were represented in 2020-2022.

**Fig. 1.**
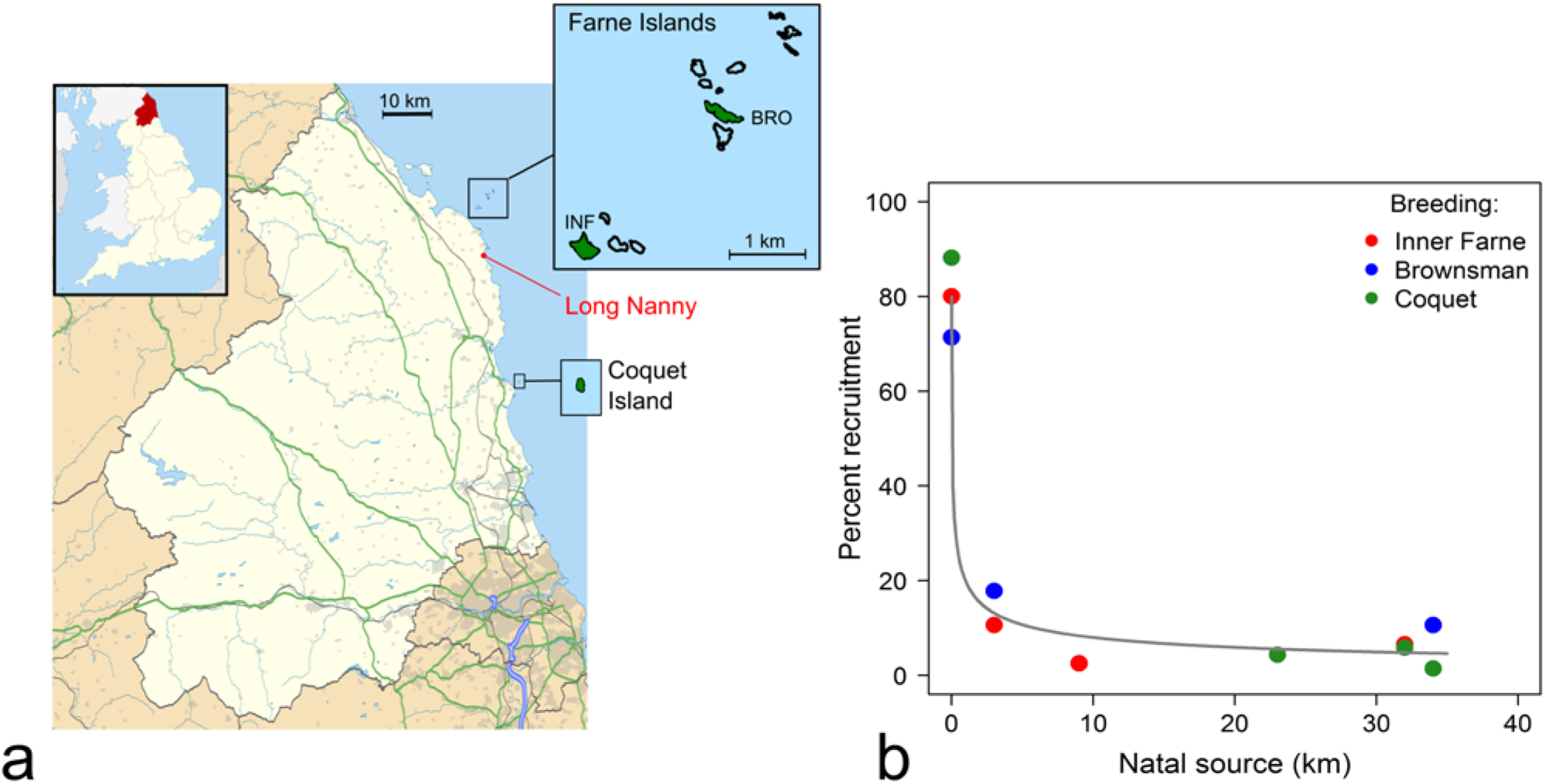
Study locations with Arctic Tern breeding colonies and recruitment in relation to distance. **a**, Map of the UK indicating the locations of Farne Islands, with breeding colonies on Inner Farne (INF) and Brownsman (BRO), Coquet Island, and the intermediate mainland site at the Long Nanny (red lines and text). Nestlings were ringed at all four sites, but nesting adult Arctic Terns were only trapped for ringing on Inner Farne, Brownsman and Coquet Island. Modified from https://commons.wikimedia.org/w/index.php?curid=11604730 with Boundary-Line™ [SHAPE geospatial data], Scale 1:10000, Tiles: GB, Updated: 10 September 2024, Ordnance Survey (GB), Using: EDINA Digimap Ordnance Survey Service, <https://digimap.edina.ac.uk>, Downloaded: 2024-11-15. **b**, Recruitment in relation to distance from the colony within Northumberland: estimated percentage composition (capture totals adjusted by numbers of nestlings ringed in natal colonies is plotted against distance of natal origin from the breeding colony. Point colours represent the three colonies at which known-age breeding adults were captured. Natal origin were the three breeding colonies and the Long Nanny colony. The grey line is a three-parameter exponential curve fitted to the data describing the relationship between percentage recruitment and distance to natal source (Taylor 1980).

From the start of the study until the end of the 2019 breeding season, the colony on Inner Farne where most trapping took place was relatively stable in number of breeding pairs; the Brownsman colony was more variable and started to decline from 2011 and the colony on Coquet Island has increased gradually by about 20 pairs per year from 2000 to 2021 (Redfern & Steel 2024). From the 2020 breeding season, unexpected consequences of the COVID 19 pandemic resulted in a catastrophic decline in terns nesting on Inner Farne, with practically complete colony abandonment in 2021, and changes in the nesting distribution of Arctic Terns on Brownsman; this was a consequence of the absence of staff and visitors making the colonies more susceptible to predation by large gulls (Redfern & Steel 2024).

Birds were trapped from mid-May to mid-July (see Results text), generally on nests with eggs but occasionally with young chicks. Nest contents at the time of capture (number of eggs and/or chicks) were recorded on most trapping occasions. On the Farne Islands, adult Arctic Terns of unknown nesting status were also caught for ringing or retrapped using untargeted methods (mainly mist nets, hand nets). Biometric data were recorded for all captures of adult birds but only head length, measured with stopped callipers (Green 1980) from the back of the skull to the bill tip, was used here; however, head length was not measured on some birds caught before 2003. Of the head-length measurements, 62% were by authors CR and DS (13:1 CR/DS), 17% by experienced ringers trained by CR and the rest were mostly done and checked under direct supervision of CR. Birds were not sexed. All trapping and ringing was licensed by BTO on behalf of UK government.

### Data and analysis

Data structuring and statistical analyses were done using R, version 4.2.0 and 4.2.2 (R Core Team 2019). Most analyses were for the first capture event of adults caught on the nest and which had been ringed previously as nestlings, mainly in the breeding colony or from other Northumberland colonies. The main analysis was focused on natal recruitment and natal dispersal between Northumberland colonies from 1995 to 2019. No Arctic Tern nestlings or adults were ringed on the Farne Islands from 1987 to 1995 inclusive; a small number of birds ringed prior to 1987 from Northumberland colonies or elsewhere were excluded to limit the main analysis to the period 1995 to 2019 for which detailed data were available; there were no significant events resulting in marked colony instability during this period. However, Arctic Terns were trapped on Coquet Island in 2020-2022 during a period of instability in the Arctic Tern colonies on the Farne Islands resulting from COVID-19 lockdown (Redfern & Steel 2024); known-age birds from this Coquet Island cohort were analysed separately. To compare natal recruitment between sites during the period 1995 to 2019, the number of captures of known-age birds originating from different sites within Northumberland was corrected for the number of nestlings ringed in previous years (up to two years before the last trapping dates to account for the age at which Arctic Terns start to breed) at different natal sites.

Although monomorphic in plumage, male terns are usually the larger sex and by discriminant analysis head length is the best single morphometric for sexing (Fletcher & Hamer 2003, Devlin *et al*. 2004, Ledwoń 2011, Monticelli *et al*. 2024). Fletcher and Hamer (Fletcher & Hamer 2003) derived means and standard deviations for head lengths of Arctic Terns nesting on Coquet Island and sexed using behavioural criteria; sexing on behaviour was completely concordant with molecular sexing for a sample of birds (Fletcher 2002). Assigning sex on a biometric criterion to mixtures of male and females is a problem of conditional probability (Bayes’ Theorem) dependent on the means and variances for the biometric in each sex, and the likely proportion of each sex in the mixture. Therefore, the means and standard deviations (sd) for head length from Fletcher and Hamer (2003) (female 70.2 mm, sd 2.51; male 72.7 mm, sd 1.68) were used to fit two-component mixture models using the *normalmixEM* function of the *mixtools* R package (Benaglia *et al*. 2009) to head-length data to estimate the proportions of females and males in samples of trapped birds of known age. Where relevant, the likely sex of possible outliers was obtained from the Bayesian posterior probabilities generated by the *normalmixEM* function. Few known-age birds from colonies outside Northumberland were captured and were not included in analyses to avoid bias from potential regional variation in size (*cf*. Monticelli *et al*. 2024). Note that the simplest biological interpretation of a mixture depends on the assumption that the females and males in the mixture were drawn without bias on size from the available populations of males and of females; this assumption fails if there is size-dependent behaviour (movement/dispersal or age of first breeding for example) for either sex.

For individuals retrapped as adults on more than one occasion, the mean of head length measurements were used, and each bird is represented only once in analyses. There was no age-related change in head length for retrapped individuals (GLMM on head length by age in years with individual as random effect, coefficient for age as fixed effect P > 0.7 and the model was not significantly different from the null model, ANOVA P > 0.7; Supplementary Information, Section 2). Confidence intervals (95 percentile) for female proportions were estimated by non-parametric bootstrap with the R package *boot* (Canty & Ripley 2024). Welch’s t-tests were used for comparisons of mean head lengths between samples, but the *emmeans* R package (Lenth 2023) was used for paired contrasts of mean head length at consecutive ages (age was coded as an ordered factor) using the multivariate t method for multiple-test correction. Days in the year for the first capture events for known-age birds were analysed to ask whether young breeding birds were caught later in the season. For each age group, median capture day was calculated and 95% confidence intervals estimated by non-parametric bootstrap using *boot*. Where relevant, general additive models were fitted using the R package *mgcv*.

## RESULTS

### Recruitment to Arctic Tern colonies - nesting birds of known origin during a period of stability

Within Northumberland, breeding dispersal of Arctic Terns is generally low (Redfern & Steel 2024). Up to 2019, the majority of adult birds of known age (ringed as nestlings) were nesting in their natal colony; the rest were mainly from other colonies in Northumberland but four of the 286 known-age birds originated from colonies outside Northumberland, two from the Isle of May and two from the Port of Dublin (Table 1). Excluding these four birds, apparent natal philopatry was 71% to 88% (Table 1), with no significant differences between colonies (Fisher’s Exact tests, P > 0.1; Table 1). Recruitment origin declined steeply with distance, and non-natal recruitment showed little variation with distance from the breeding colonies (Fig. 1b). Nearly two-thirds of the data for known-age birds were based on trapping results from Inner Farne, with ∼ one-tenth from Brownsman and a quarter from Coquet Island (Table 1).

### Sex bias in local dispersal and recruitment

In terns, males generally have longer heads (head + bill) and this is the best single morphometric for sexing with accuracies of 72% or more (Fletcher & Hamer 2003, Devlin *et al*. 2004, Ledwoń 2011, Monticelli *et al*. 2024). The mean head length of non-natal nesting birds was significantly shorter than birds nesting on their natal site (Welch two-sample t. test; t = −2.63, df = 52.8, P = 0.01; non-natal mean 70.52 mm, sd = 2.37, n = 39; natal mean 71.6 mm, sd = 2.33, n = 207). For all birds of known origin with head-length data (natal and non-natal groups combined; n= 246), the distribution of head lengths was consistent with proportions of 0.55 female to 0.45 male (female proportion 95-percentile confidence range 0.42 to 0.67), and was similar to all data for nesting Arctic Terns caught across the three Northumberland sites (Fig. 2a, n = 1217 individuals; female proportion 0.55, 95-percentile confidence range 0.49 to 0.61). For the natal recruits, the proportion of females was 0.46 (Fig. 2b; 95 percentile confidence range 0.33 to 0.59). Conversely, for non-natal recruits, the head-length distribution was consistent with at least the majority of these birds being female (Fig. 2c; female proportion, 95 percentile confidence range 0.74 to 1). A simple interpretation of these data is that up to nearly a third of known-age breeding females were non-natal in origin whereas most males originated from the natal site.

**Fig. 2.**
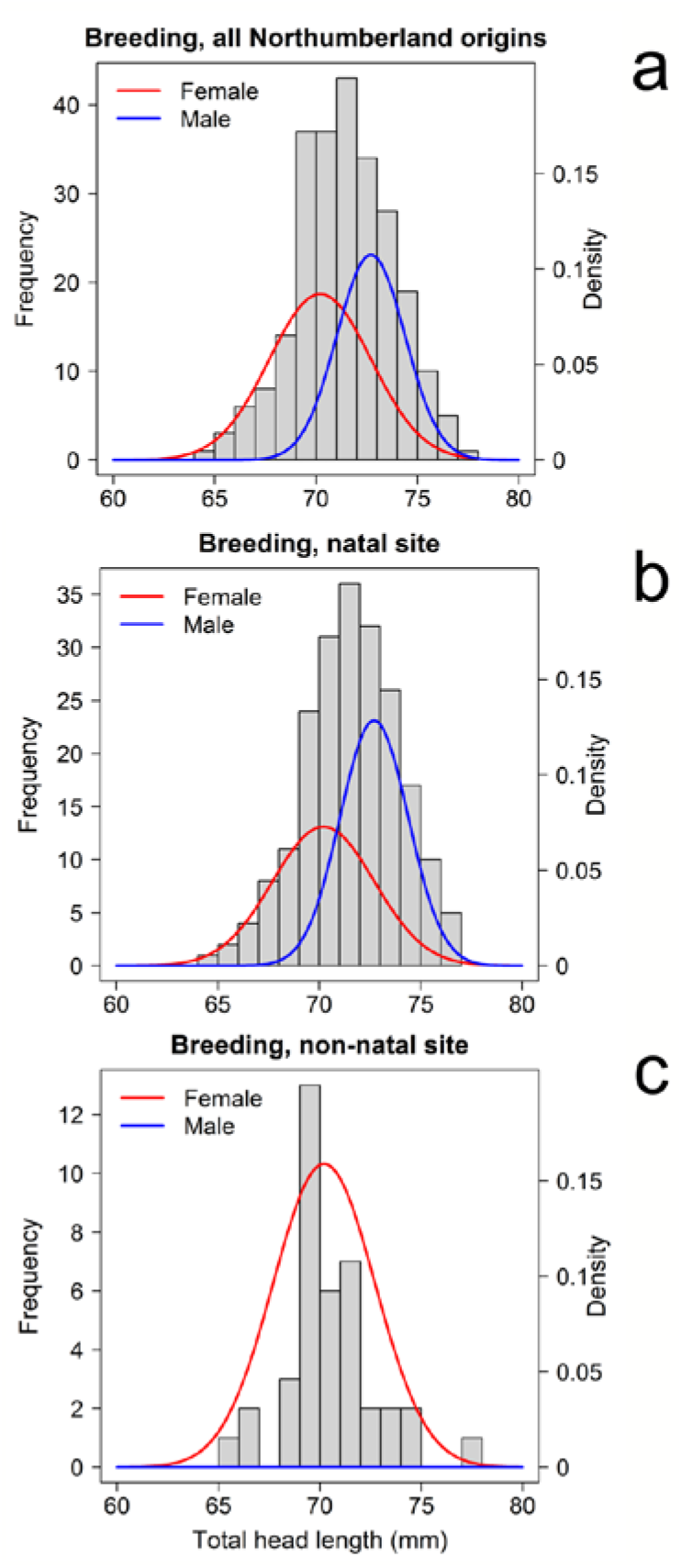
Distributions of head lengths for nesting Arctic Terns. **a**, Histogram of all data for birds trapped or retrapped as nesting adults (known-age and unknown age) with normal curves for head lengths based on means and standard deviations of nesting Arctic Terns sexed as female (red) or male (blue) (Fletcher and Hamer, 2003). Areas under curves represent the calculated mixture proportions of females (0.55) and males (0.45) from head-length data, estimated using the *normalmixEM* function from the R package *mixtools* with means and standard deviations (in parentheses) constrained to values given by Fletcher and Hamer (2003): females 70.2 mm (2.51); males 72.7 mm (1.68). **b**, Head-length distributions for natal birds, curve details as in **a**; mixture proportions: females 0.43, males 0.57. **c**, Head length distributions for non-natal birds, curve details as in **a**; mixture proportions: males < 0.0001, females 1. In this non-natal group, one bird has a head length of 77 mm. From the mean and sd for females, around 1 in 100 could be expected of this size; at the best-fit mixture proportion implying a significant female bias in natal dispersal, the probability that this bird is female is > 0.9.

### Age and sex at first breeding

From recaptures of nesting birds ringed as nestlings, Arctic Terns start to breed from two years (3^rd^ calendar year) of age (Redfern 2021, Redfern & Steel 2024). A previous study on Inner Farne, based on a smaller year-range 1966-1968 and pool of potential recruits (Supplementary Information, Section 3), did not find Arctic Terns breeding at two years of age (Horobin 1971, Coulson & Horobin 1976). However, in the present dataset, of 245 known-age adults caught on a nest and for which nest contents were recorded across all three Northumberland sites, 10 were two years old and were caught on nests with clutch sizes of one to three eggs (Supplementary Information, Section 4). For nesting Arctic Terns aged 12 years or less with head-length measurements (n = 240, sample size per age category range 11 to 36) there was significant variation in head length with age (ANOVA, type III, F10, _229_ = 2.8136, P = 0.0026), and birds aged four or less had shorter head lengths than those five years or older (Fig. 3a; contrasts on consecutive pairs). For the full dataset (n = 273), mean head length for birds aged four or less (n = 77; mean = 70.4, sd = 2.25) was significantly shorter than older birds (n = 196; mean = 71.8, sd = 2.34; t = −4.6374, df = 144.36, P < 0.00001). The data distributions suggest that the majority of the birds aged four or less (young age group) were female (95 percentile range for proportion: 0.8 to 1; Fig. 3b, c), whereas older birds were mixtures of females and males in roughly equal proportions (Fig. 3b,d). If the capture of non-natal breeders was biased towards young females, then this could affect interpretation of the head-length data for natal versus non-natal birds. However, only a quarter of the non-natal birds were aged 4 or less (Supplementary Information, Section 5) and, therefore, age was not likely to be a factor in the female bias for dispersal from the natal colony.

**Fig. 3.**
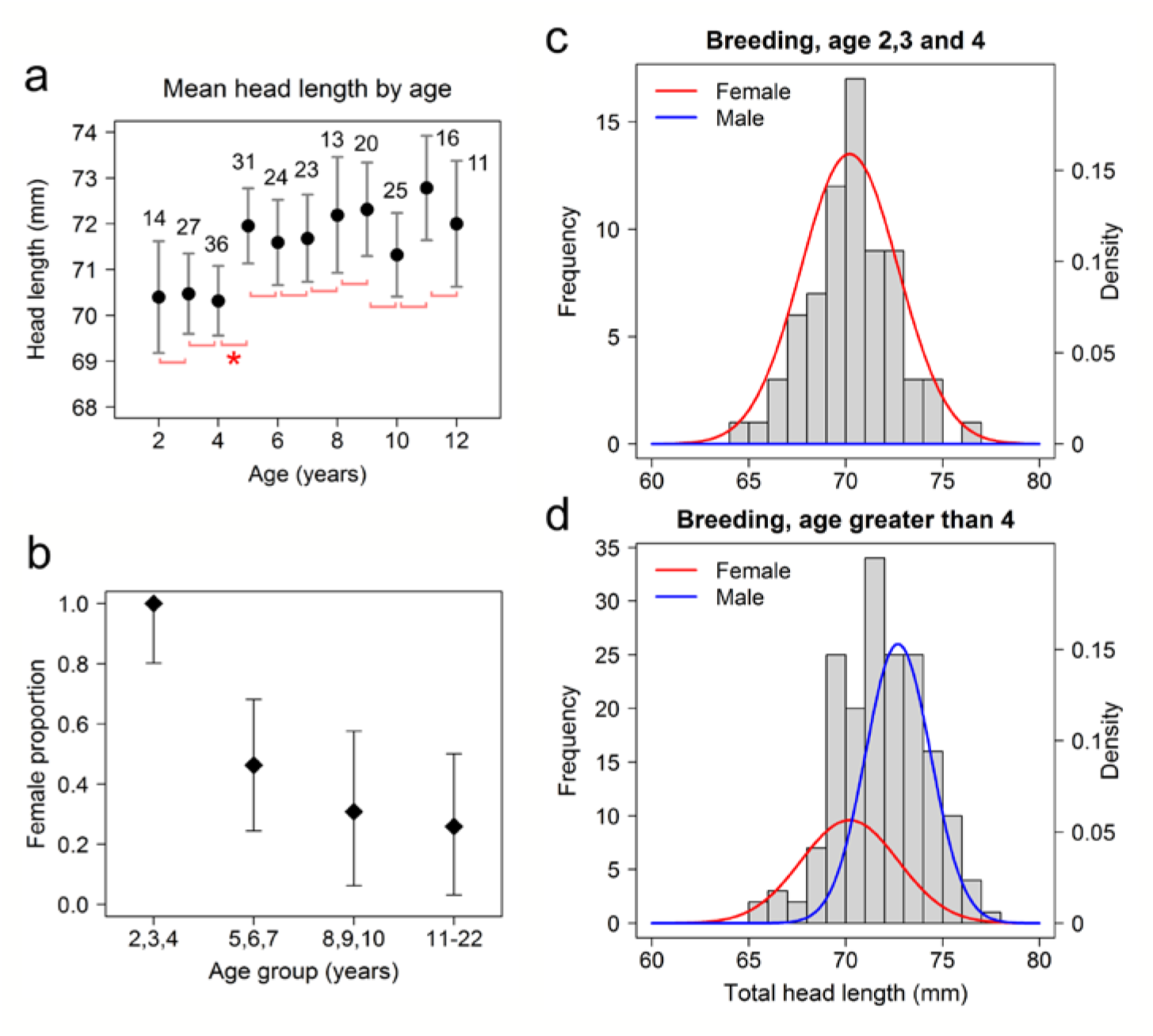
Head-length distributions of known-age nesting birds. **a**, mean head length by age up to 12 years old; grey error bars are 95% confidence intervals. Horizontal pink square brackets at the ends of lower confidence limits indicate consecutive (ordered by age) pairwise comparisons of head length; all were non-significant except the age 4 to age 5 comparison indicated by the red asterisk (P = 0.04). Sample sizes are given over the upper 95% confidence limits. **b**, female proportion in three-year age groups; error bars are 95 percentile ranges from bootstrapping. **c**, frequency distribution of head lengths for the young age group (2-4 years old inclusive) and fitted mixture proportions (female proportion and 95-percentile range in b). **d**, frequency distribution of head lengths for all older birds (greater than 4 years old) and fitted mixture proportions. For all older-age birds, the female proportion was 0.36 (95-percentile range 0.22 to 0.49).

Birds in the young age group, particularly the two-year-old birds, tended to be caught later in the season compared to birds of five years or more in age (Fig. 4a). The distribution of catching effort (Fig. 4b) suggests that young birds were not likely to be on nests earlier in the season. All except two of the 77 birds that were four years old or younger were on nests with eggs or young chicks at the time of capture and clutch/brood sizes were not significantly different from older birds (Fisher’s exact test, P = 0.45; Supplementary Information, Section 4). The median times of capture for age-two and age-three birds differed from those of older birds by similar magnitudes to the times of arrival of these age classes in Jean Horobin’s study on Inner Farne in 1966 – 1968 (Supplementary Information, Section 3; Horobin 1971, Coulson & Horobin 1976).

**Fig. 4.**
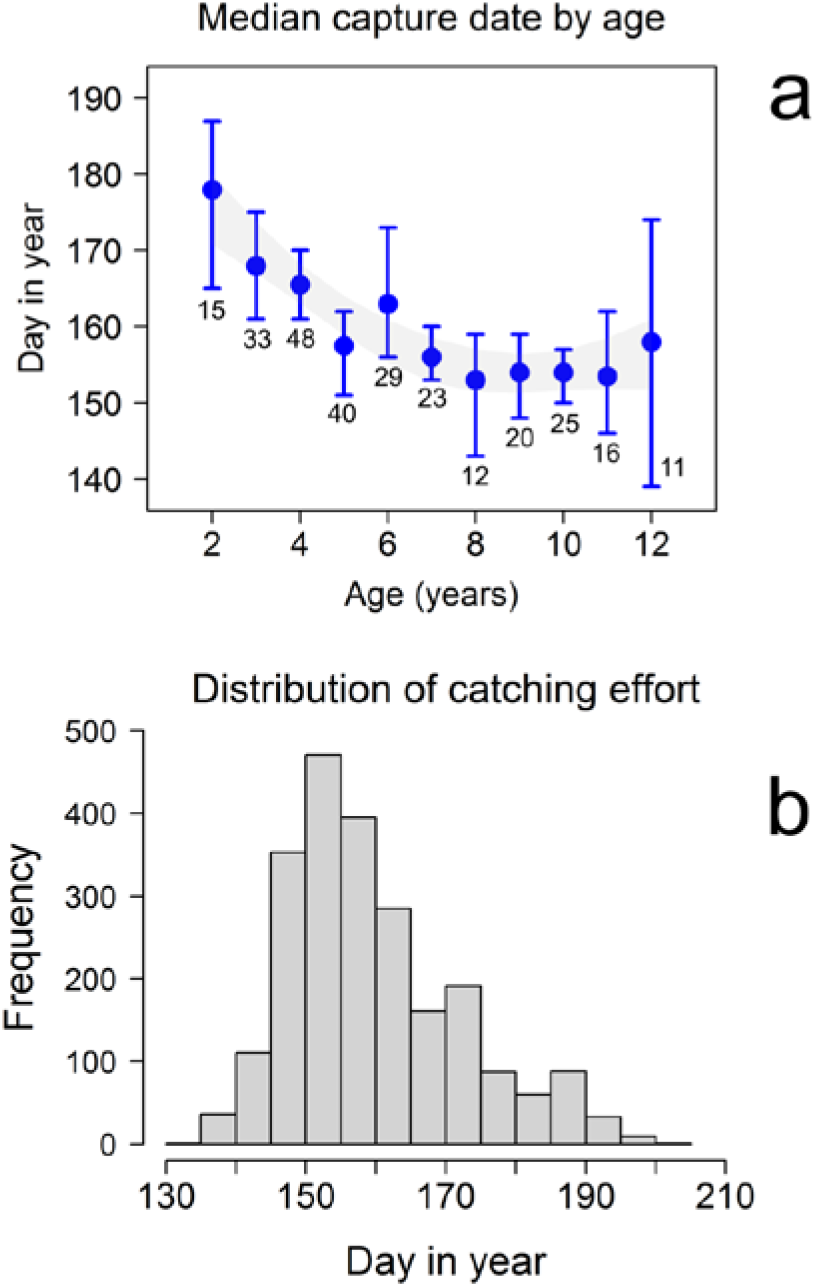
Capture dates of known-age birds. **a**, median capture date by age for birds up to age 12 years; error bars are 95% confidence intervals by non-parametric bootstrap and the shaded grey area is the 95% confidence range for medians from a general additive model. Numbers below the error bars are sample sizes for each age. **b**, distribution of catching effort (total birds caught on the nest) for all nesting birds for the study period overall.

These data indicating a sex-specific bias in age at first breeding raise the question whether or not males of similar ages are present in the colony but not breeding, or less likely to be captured on the nest. To test this, the head length distributions and female proportion were analysed for known-age birds caught other than on nests within the Inner Farne and Brownsman colonies during the breeding season. For these birds, the head-length distribution (Supplementary Information, Section 6) and female proportion for birds aged two to four inclusive (n = 34) were similar to breeding birds caught on the nest (1, 95 percentile range 0.65 to 1), but mean head length was not significantly shorter than older birds (n = 64) not caught on the nest (t - 1.34, df = 65.4, P = 0.18; mean for age < 5, 70.1 mm, sd = 2.63; mean for ages > 4, 70.9 mm, sd = 2.55). For these older birds, the female proportion was 0.77 (95-percentile range 0.53 to 1; Supplementary Information, Section 6). The mean head length for birds of non-natal origin (n = 30) was significantly shorter than those of natal origin (n = 71; t = 3.02, df = 46.3, P = 0.004; mean for non-natal, 69.4 mm, sd = 2.51; mean natal, 71.1 mm, sd = 2.47), and the head length distributions (Supplementary Information, Section 6) and proportion of females (natal 0.65, 95-percentile range 0.41 to 0.88; non-natal 1, 95-percentile range 0.98 to 1) were similar to known-age birds caught on the nest. These results suggest that young males and non-natal males were less likely to be present within the colony compared to young or non-natal females.

### Recruitment to Coquet Island after 2019

After 2019, Inner Farne was largely abandoned by Arctic Terns as a breeding colony from 2020 to 2021 (Redfern & Steel 2024). COVID-19 lockdown prevented further work on the Farne Islands, but some nest trapping was able to continue on Coquet Island in the period 2020 to 2022. The pattern of apparent natal recruitment (recruitment of birds ringed as nestlings) to Coquet Island in 2020 – 2022 was significantly different to the period up to 2015 (Fisher’s exact test, P < 0.00001; Table 1): in 2020 – 2022, of 27 nesting adults of known natal origin, 18 (67%) were from Inner Farne and Brownsman (Table 1). Contrary to known-age nesting birds recaptured before 2020, there was no significant difference in head length between the non-natal and natal birds (t = −0.131, df = 14.726, P = 0.9, non-natal mean 71.2 mm, sd = 2.72; natal mean 71.3 mm, sd = 3.01), and the distribution of head lengths suggests similar proportions of females within the non-natal and natal groups (non-natal: 0.72, 95-percentile range 0.234 to 1; natal: 0.7, 95-percentile range 0.29 to 1; Fig. 5). The age distribution of natal birds ranged from 5 to 22, and non-natal birds from 3 to 19 (Supplementary Information, Section 7). The sample size for birds age 4 or less is small (n = 5), and these were all non-natal in origin with no significant difference in head length from older birds (t = 0.08, df = 6.1203, P = 0.94; mean age < 5, 71.3 mm, sd = 2.74; mean age > 4, 71.2 mm, sd = 2.84), consistent with a composition of females and males for both age groups (Supplementary Information, Section 8). These data suggest that non-natal recruits to Coquet Island in 2020 – 2022 mainly represented breeding dispersal as a consequence of colony abandonment on the Farne Islands in 2020 – 2021.

**Fig. 5.**
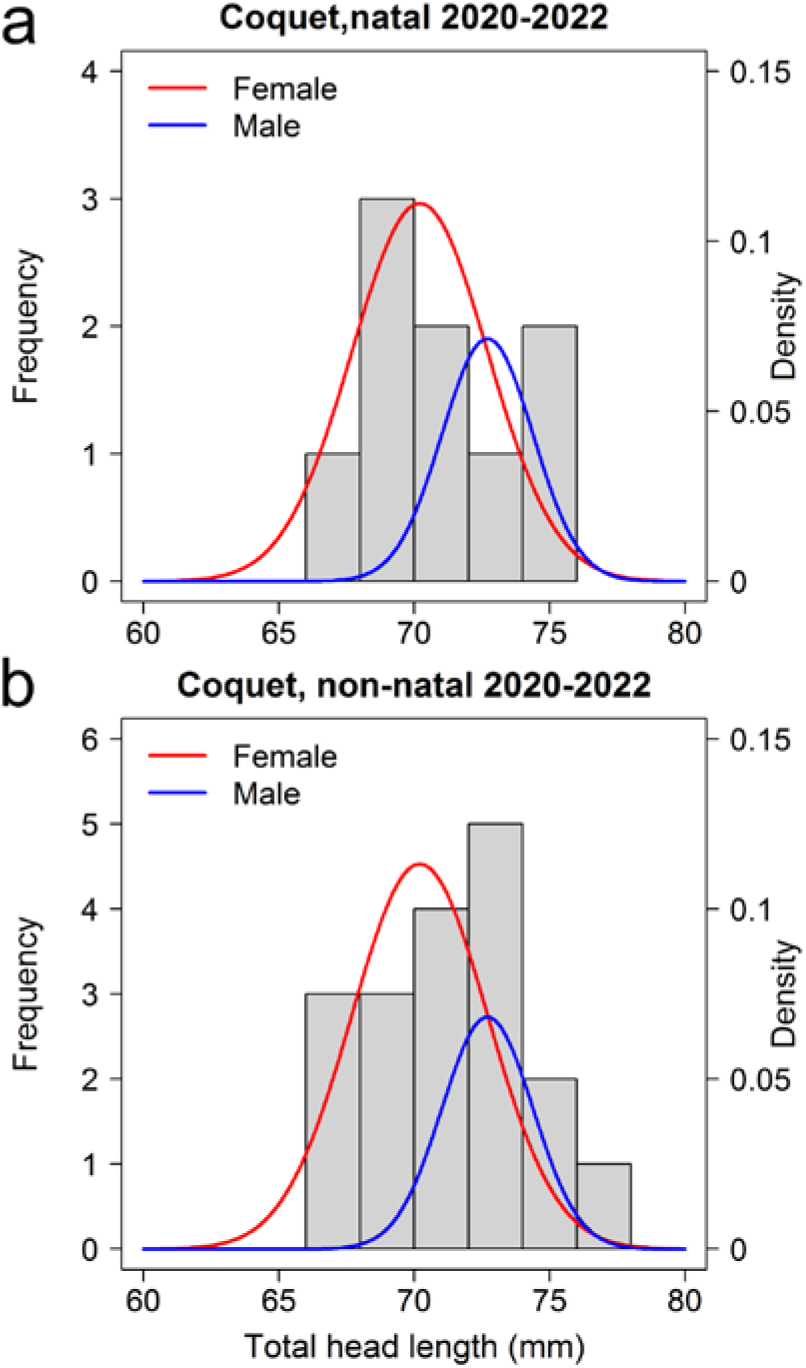
Distributions of head lengths for Arctic Terns nesting on Coquet Island in 2020 - 2022. **a**, Histogram of all data for natal-origin birds with normal curves for head lengths based on means and standard deviations of sexed nesting Arctic Terns (Fletcher and Hamer, 2003). Details as in Fig. 2. **b**, Head-length distributions for all nesting birds of non-natal origin, curve details as in **a**.

## DISCUSSION

### Natal philopatry

Nesting birds caught on the nest are, by definition, breeding members of the colony. With respect to age and natal/non-natal status, the head-length distributions of known-age birds not caught on the nest were similar to nesting birds, implying that they could be regarded as *bona fide* colony members. Nevertheless, to avoid ambiguity, we limit the main analysis to birds known to be breeding. Determining the extent of natal philopatry from recaptures of ringed birds can be difficult because birds moving elsewhere may be missed and the origins of unringed birds are unknown (Coulson 2016). However, we can be reasonably confident about the extent of natal philopatry because of the extensive ringing of nestlings in Northumberland colonies for many years. At the three island sites, natal philopatry was high with a relatively low recruitment from other Northumberland colonies. Arctic Terns have one of the longest annual migrations of any bird species and in this context the degree of natal philopatry is remarkable. Navigational techniques used by Arctic Terns, particularly those from North-west European populations with their extensive crossing of the southern Indian Ocean (Redfern & Bevan 2022), are likely to include magnetoreception (Mouritsen 2018) which may also facilitate natal philopatry as a consequence of imprinting to magnetic declination at natal sites (Wynn *et al*. 2020).

The few birds known to have originated from outside Northumberland came from colonies around the Irish Sea or further north on the North Sea coast. These colonies account for around 36% of the British Isles population (Wyn 2023). The Irish Sea may be a major migration route of Arctic Terns to the Farnes (Redfern and Bevan 2020), facilitating the recruitment of new breeders to the east coast of the UK from Irish Sea colonies. Migration routes of Arctic Terns nesting further north and west in Scotland and Ireland are unknown. Without an understanding of the extent to which migration routes facilitate recruitment of new breeders from different parts of the North-west European range, it is difficult to assess the likely population size from which new recruits to Northumberland colonies are derived. Given the relationship between recruitment and distance to non-natal colonies, the extent of recruitment from outside Northumberland is likely to be relatively small.

For Inner Farne, the extent of natal philopatry was similar to Brownsman and Coquet Island; these should be regarded as separate colonies within a metapopulation (*contra* Antaky et al. 2021). Natal philopatry could be an indicator of habitat quality (Antaky *et al*. 2021, Byer & Reid 2022), perceived by fledglings (Serrano *et al*. 2021) or by returning pre-breeders prospecting for breeding sites (Crespin *et al*. 2006). All three islands on which nest trapping took place have small numbers of resident staff during the breeding season, but only Inner Farne is exposed to high numbers of visiting public each day during the breeding season (Redfern & Steel 2024). Most Arctic Tern chicks on Inner Farne would develop in an environment where high exposure to human activity is normal and from conspecific social interactions might perceive it as a habitat of good quality; this implies that high levels of human activity do not discourage fledglings from returning as potential breeders. Furthermore, this also implies that terns from other sites without the same prior social context, both human and conspecific, are not discouraged by human activity from settling to breed.

### Sex ratio and catchability

Confidence intervals for estimates of the proportion of females and males from head-length data for all nesting birds were biased towards a slightly greater proportion of females. These are dependent on the sample distributions for Coquet females and males sexed from behavioural observations (Fletcher & Hamer 2003), but also could reflect differential catchability of females and male birds (McKnight & Ligon 2017). For Arctic Terns, both sexes share incubation (Cramp 1985), but not necessarily equally. Horobin (1971) reported that on Inner Farne in the late 1960s females were found incubating on 54% of occasions. That would fit with the slightly higher proportion of females to males in the nesting birds sampled here. For Roseate Terns, a similar slightly higher female proportion (56%) was reported to be a result of female-female pairs (Nisbet & Hatch 1999). If true for Arctic Terns, then the assumption of a completely monogamous breeding system might not strictly hold, or could be age-dependent; paternity and molecular sexing studies to address such questions would be useful.

### Sex bias in dispersal

Head-length data show that natal dispersal is biased towards birds with short head lengths. The inference from mixture models constrained by means and standard deviations of sexed Arctic Terns nesting on Coquet Island (Fletcher & Hamer 2003) is that non-natal birds were predominantly female and represented up to around a third of known-age females. The corollary is that male natal philopatry is very high, with most of the reduction in natal philopatry to the observed 71%-88% accounted for by the natal dispersal of females. This female bias in dispersal to non-natal sites within the Northumberland metapopulation fits with the general expectation for bird species (Greenwood 1980, Greenwood & Harvey 1982, Clarke *et al*. 1997, Li & Kokko 2019). Offspring and parents conflict in their preferences for offspring dispersal (Roff 1975, Hamilton & May 1977), but costs from dispersal can be mitigated by the reduction in potential for inbreeding (Szulkin & Sheldon 2008). Dispersal may also vary in relation to population or habitat stability (Hamilton & May 1977, Spicer & Forys 2023), noting the behavioural distinction between dispersal and dispersal distance (Liberg & von Schantz 1985, Mabry *et al*. 2013).

Female-biased natal dispersal is thought to be a consequence of social monogamy (Greenwood 1980, Greenwood and Harvey 1982, Trochet et al. 2016, Li and Kokko 2019) but this is not supported when phylogenetic relationships are taken into account (Mabry *et al*. 2013) and the mechanisms are unclear because consequences of dispersal in terms of genetic fitness can be hard to measure in natural populations. While the conclusion that natal dispersal of Arctic Terns is female biased in Northumberland seems robust, other contributors to the short head lengths of non-natal birds should be considered. For example, it is possible that small males might be included in the non-natal birds if size-dependent dispersal of small males was a behavioural strategy related to sexual-size dimorphism in Arctic Terns or a consequence of competitive exclusion by larger males on natal sites. Further studies are needed to test this possibility.

### Age of first breeding and bias by size and sex

Age at first breeding is a demographic parameter that can vary by environmental constraints and individual quality (Fay *et al*. 2016). Thus, the difference between the present data for Northumberland Arctic Terns and the earlier study (Horobin 1971, Coulson & Horobin 1976) with respect to birds breeding at two years old is probably not of significant interest. Nevertheless, breeding birds of age two years can make a contribution to population demography, albeit small. Given the extensive migration of Arctic Terns, it is not surprising that individuals would attempt to breed as early as possible to enhance life-time reproductive success.

As with natal dispersal, head-length distributions suggest that females breed up to three years earlier than most males, with the first females breeding at two years of age. A bias towards earlier breeding ages for females has also been reported for Common Tern *Sterna Hirundo* (Becker *et al*. 2008). Conversely, Horobin (Horobin 1971, Coulson & Horobin 1976) found that of nine Arctic Tern pairs, sexed by behaviour, where the females were age three or four, six of the males were of similar age. Although sample size and environmental factors such as body condition and colony density or stability may result in annual variation in the ages of first breeding by males and females, there are biological factors that might result in sex-specific optimal breeding ages for Arctic Terns.

The apparent bias towards females in birds aged four or less is not likely to be an issue of catchability because similar results were obtained with Arctic Terns caught using methods that did not directly target nesting birds. This then raises the question whether females have more to gain than males by breeding early. The higher mortality cost to females from egg production (Romano *et al*. 2022) and potential costs of natal dispersal (Payevsky 2016), could be mitigated by breeding at an earlier age than males. An additional strategy that young females could use to enhance reproductive success would be to mate with suitable available males, either in the colony or at colony roost sites, and lay eggs in the nests of older females. Conspecific brood parasitism is closely related to coloniality (Yom-Tov 2001), has been reported for Roseate (Shealer & Zurovchak 1995) and Common Terns (Rohwer & Freeman 1989), and in other contexts can be a strategy used by young, inexperienced females (Lyon & Eadie 2008). This is an understudied aspect of tern breeding biology important for a full understanding of their population demography.

With respect to males, slightly longer head length implies sexual selection with larger males having a competitive advantage (Caron & Pie 2024). Phylogenetically-controlled analyses confirm hypotheses that sexual selection on male size in birds is associated with breeding at an older age (delayed maturation; Ancona *et al*. 2020). Although Arctic Terns are assumed to have a socially monogamous mating system (Hatch *et al*. 2020), the sexual size dimorphism implies a degree of polygamy (extra-pair paternity) outside the pair bond (Valcu *et al*. 2023), a behaviour also associated with delayed maturation (Ancona *et al*. 2020). Thus, male Arctic Terns might be expected to breed at an older age than females. Overall, there are good biological explanations for both females and males to breed at different ages.

There is one speculative caveat that cannot be excluded without further work and that is the question whether smaller males (less delayed maturation with respect to developmental pathways) could compensate for size by breeding earlier than larger males. In other sexually dimorphic animals, strategies to find mates can vary according to size (Carroll & Salamon 1995). Given evidence for inherited characteristics for dispersal in some bird species (Korsten *et al*. 2013, Payevsky 2016), we should also be open to the possibility that the timing of breeding of males could be associated with genetic determinants of head length.

### Natal versus breeding dispersal

Natal dispersal, the movement of birds from a natal site to a different first-breeding site, is distinct (behaviourally at least) from breeding dispersal, the movement of breeding birds from one location or colony to another (Greenwood & Harvey 1982, Cantrell *et al*. 2017). Other than long-term trends for gradual increases or decreases, Arctic Tern populations in Northumberland were relatively stable up to 2019, and instances of known breeding dispersal were rare (Redfern & Steel 2024). Captures of known-age birds (ringed as nestlings) during that period were interpreted as representing natal philopatry and natal dispersal. Conversely, head-length comparisons with respect to natal/non-natal status for nesting birds caught on Coquet Island in 2020-2022 indicate that the apparent natal recruitment was mostly breeding dispersal of birds that had abandoned Inner Farne, and possibly also Brownsman, in 2020-2021 as a result of the disruption precipitated by unavoidable withdrawal of staff and visitors from the islands in consequence of COVID-19 lockdown regulations (Redfern & Steel 2024); a few of those birds may have been first-time breeders unable to recruit to Inner Farne or Brownsman. A corollary of these events is that breeding dispersal precipitated by sudden changes in colony-site suitability could be misinterpreted in terms of natal philopatry/dispersal, particularly if those changes become fixed and/or unrecognised. In other metapopulations or contexts, colony-site suitability could fluctuate regularly and result in greater fluidity of breeding dispersal between years (Egevang & Frederiksen 2011), making natal philopatry and natal dispersal difficult to assess.

At a finer scale, it may be harder to distinguish between natal dispersal and breeding dispersal than realised. In an intensively-studied area of the Inner Farne colony, young breeding birds aged two or three years were under-represented in subsequent recapture events (Redfern & Steel 2024). Assuming this was not a consequence of higher mortality in this age group, where these younger birds ended up is unknown, but it is possible that in subsequent breeding seasons some young first-breeders, whether successful or not, could breed elsewhere within the Inner Farne colony or further afield. Strictly, this would be breeding-dispersal and could be a consequence of poor breeding outcome (Bialas *et al*. 2023) or stability of the pair bond, particularly if young returning breeders are tardier in returning to the breeding colony than a more-experienced partner (Horobin 1971, Coulson & Horobin 1976, Redfern 2021).

Detecting breeding-dispersal events of young birds will be difficult without extensive and long-term capture-recapture/resighting programs involving the ringing of adults and nestlings across the metapopulation. Breeding dispersal within or between colonies may be just a question of spatial scale where dispersal probability declines with distance in the same way as natal dispersal (Taylor 1980, Cantrell *et al*. 2017) and may be influenced by age and/or breeding success (Johst & Brandl 1999, Kim *et al*. 2007, Eeva *et al*. 2008), as well as landscape features (Sieger & Hovestadt 2021). Arctic Tern colonies in Northumberland have benefited by active conservation for three decades or more. Local events such as the disruption of forage-food availability by a population explosion of Snake Pipefish *Entelurus aequoreus* (Harris *et al*. 2007) and potential botulism (Redfern *et al*. 2020) have been short-term constraints on the numbers and productivity of breeding adults. Disruption to conservation management by COVID-19 (Redfern & Steel 2024) and the effects of HPAI may prove to be more of a problem, and at a broader scale in Britain and Ireland Arctic Terns are now of substantial conservation concern (Gilbert *et al*. 2021, Stanbury *et al*. 2024). Sex-biased dispersal, age of breeding and the links between colonies offer new perspectives on Arctic Tern population biology which will benefit future conservation planning.

## ACKNOWLEDGEMENTS

We thank Mike Hodgson for details of his Arctic Tern chick ringing in the Long Nanny colony, Brian Burke for data from the Port of Dublin, Anne Wilson for Northumberland seabird monitoring data and the Natural History Society of Northumbria for support. We are especially grateful to the National Trust Rangers who have collected and supplied data for counts of Arctic Tern nesting pairs on the Farne Islands over many years, and for their interest and help with the study. Finally, CR thanks Dr Natasha Murphy for her question on natal philopatry at the end of a presentation on Arctic Terns at the Welsh Ringing Course which stimulated some of these analyses.

## Contributor roles

CR: conceptualisation, data curation, investigation, methodology, formal analysis, wrote the original draft, review and editing of the manuscript. DS: Provision of resources, investigation, review and editing of the manuscript. PM: Provision of resources, investigation, review and editing of the manuscript.

