## Supplementary Information for "Natal philopatry, dispersal and age of first breeding in relation to size and sex of Arctic Terns *Sterna paradisaea*"

0000-0002-1833-8048 (CPF Redfern)

**Index**


| Section | Heading | Page |
|---|---|---|
| 1 | Nestlings ringed in Northumberland 1995 to 2019; Table S1 | 2 |
| 2 | Model statistics to test for change in head length with age | 3 |
| 3 | Coulson & Horobin 1976 study context; Table S2 | 4 |
| 4 | Clutch size by age groups of known-age birds; Tables S3, S4, S5 | 4 |
| 5 | Age distribution of natal and non-natal nesting birds; Table S6 | 5 |
| 6 | Fig. S1: Birds not caught on the nest: head length by age and natal status | 6 |
| 7 | Coquet 2020-2022: age of natal and non-natal birds; Tabel S7 | 6 |
| 8 | Fig. S2: Coquet 2020-2022: head-length distributions by age class | 7 |
| | REFERENCES | 8 |

*SECTION 1*

**Table S1:** Arctic Tern nestlings ringed in Northumberland 1995 to 2019. For 1995 to 2019 (all year ranges are inclusive), Arctic Tern nestlings ringed in Northumberland represented 26.7% of the total for Britain and Ireland over the same period¶.

| Year | Inner Farne | Brownsman | Coquet Island | Long Nanny* | Total | %Britain & Ireland¶ |
|---|---|---|---|---|---|---|
| **1995** | 0 | 0 | 197 | 37 | 234 | 3.6 |
| **1996** | 294 | 113 | 222 | 115 | 744 | 17.7 |
| **1997** | 263 | 185 | 189 | 115 | 752 | 12.7 |
| **1998** | 228 | 107 | 176 | 114 | 625 | 13.1 |
| **1999** | 110 | 12 | 212 | 128 | 462 | 10.7 |
| **2000** | 401 | 204 | 311 | 383 | 1299 | 23.7 |
| **2001** | 185 | 0 | 244 | 363 | 792 | 25.6 |
| **2002** | 114 | 28 | 300 | 190 | 632 | 15 |
| **2003** | 111 | 89 | 254 | 194 | 648 | 15.9 |
| **2004** | 69 | 40 | 144 | 282 | 535 | 28.8 |
| **2005** | 127 | 39 | 152 | 244 | 562 | 16.6 |
| **2006** | 145 | 111 | 142 | 211 | 609 | 16.2 |
| **2007** | 298 | 164 | 219 | 51 | 732 | 38.8 |
| **2008** | 111 | 251 | 98 | 106 | 566 | 44.1 |
| **2009** | 341 | 127 | 207 | 54 | 729 | 25.9 |
| **2010** | 195 | 264 | 109 | 0 | 568 | 47.5 |
| **2011** | 208 | 183 | 151 | 284 | 826 | 45.8 |
| **2012** | 545 | 311 | 66 | 155 | 1077 | 46 |
| **2013** | 726 | 453 | 96 | 231 | 1506 | 46.4 |
| **2014** | 501 | 537 | 92 | 235 | 1365 | 35.8 |
| **2015** | 318 | 220 | 92 | 36 | 666 | 29.5 |
| **2016** | 494 | 115 | 104 | 9 | 722 | 27 |
| **2017** | 523 | 16 | 86 | 130 | 755 | 33.3 |
| **2018** | 193 | 134 | 65 | 235 | 627 | 27.5 |
| **2019** | 187 | 149 | 82 | 45 | 463 | 21.1 |

*Long Nanny data from Mike Hodgson

¶Calculated from ringing totals in BTO Ringing Report (Robinson *et al.* 2024). A small colony of Arctic Terns with Common Terns *Sterna hirundo* on or near Holy Island (Lindisfarne) is excluded from Northumberland totals. The numbers of breeding pairs there are small (Dean *et al.* 2015) and often only monitored as a total for both species (recorded as 'Commic' terns). We are not aware that any terns have been ringed there during the period of this study.

*SECTION 2*

**Model statistics to test for change in head length with age**

The glmmTMB package in R (Brooks *et al.* 2019) was used to test whether head length changed by age. Bird identity (ring number) was a random effect and age in years as the fixed effect: age was non-significant (P > 0.7) and the model (m1) was not significantly different (P > 0.7) from the null model (m2).

m1 <- glmmTMB(head length ~ age + (1| ring number), data= known-age nesting Arctic Terns)
m2 <- glmmTMB(head length ~ 1 + (1| ring number), data= known-age nesting Arctic Terns)

Summary, m1: Gaussian (identity)
Formula: head length ~ age + (1 | ring number)

| AIC | BIC | log Likelihood | deviance | residuals degrees of freedom |
|---|---|---|---|---|
| 828.3 | 842.7 | -410.2 | 820.3 | 263 |

Random effects:
Conditional model:

| Groups | Name | Variance | Standard deviation |
|---|---|---|---|
| Ring number | (Intercept) | 6.0358 | 2.4568 |
| | Residual | 0.3753 | 0.6127 |

Observations: 267, groups, ring number: 84
Dispersion estimate for Gaussian family ($sigma^2$): 0.375

Conditional model:

| | Estimate | Std. Error | z value | Pr(>\|z\|) |
|---|---|---|---|---|
| (Intercept) | 71.136796 | 0.320531 | 221.93 | <2e-16 *** |
| Age | 0.006729 | 0.018598 | 0.36 | 0.717 |

Significance codes: 0 ‘***’ 0.001 ‘**’ 0.01 ‘*’ 0.05 ‘.’ 0.1 ‘ ’ 1

ANOVA(m1, m2)
Models:
m2: head length ~ 1 + (1 | ring number), zi=~0, disp=~1
m1: head length ~ age + (1 | ring number), zi=~0, disp=~1

| | Df | AIC | BIC | log Likelihood | deviance | Chisq | Chi Df | Pr(>Chisq) |
|---|---|---|---|---|---|---|---|---|
| m2 | 3 | 826.44 | 837.20 | -410.22 | 820.44 | | | |
| m1 | 4 | 828.31 | 842.66 | -410.15 | 820.31 | 0.131 | 1 | 0.7174 |

*SECTION 3*

**Coulson & Horobin 1976 study context**
Data in Jean M Horobin's PhD thesis (Horobin 1971, Coulson & Horobin 1976) are summarised here to provide context for the present study. Horobin's work was carried out on Inner Farne in 1966, 1967 and 1968 and utilised known-age birds, all except one ringed previously as nestlings on the Farne Islands, which were recaptured or resighted on Inner Farne. The bird not ringed on the Farnes had been ringed on a small tern colony on/near Holy Island, 10 km north-west of Inner Farne (see Supplementary Information, Section 1). The last year of the Horobin & Coulson study was affected by high mortality as a result of a Dinoflagelate bloom (Coulson *et al.* 1968, Robinson 1968). The known-age Farne Islands birds were ringed using British Trust for Ornithology rings issued to the Natural History Society of Northumbria (NHSN) under the auspices of Grace Hickling. Ringing totals from NHSN records for the relevant previous years with respect to birds of likely first breeding age are in Table S2.

**Table S2:** Arctic Tern nestlings ringed on the Farne Islands before 1967

| **Ringing year** | **Farne nestlings ringed** | **Horobin and Coulson study 1966 – 1968** (Coulson & Horobin 1976) | | | | | |
|---|---|---|---|---|---|---|---|
| | | **Capture 1966-age=3** | **Capture 1967-age=3** | **Capture 1968-age=3** | **Capture 1966-age=4** | **Capture 1967-age=4** | **Capture 1968-age=4** |
| **1962** | **1824** | | | | 24 | | |
| **1963** | **2459** | 11 | | | | 27 | |
| **1964** | **2387** | | 25 | | | | 37 |
| **1965** | **1698** | | | 2 | | | |
| **1966** | **1568** | | | | | | |

For Horobin and Coulson's study, age two birds caught in 1966 – 1968 would be drawn from 5653 nestlings ringed in 1964 – 1966, and age three birds from 6544 nestlings ringed 1963 – 1965.

By comparison, the total potential pool from which age-two birds or older were derived for this present study was 15676 birds ringed as nestlings (Table S1, Northumberland total from 1998 to 2017). So, although no breeding birds of age two were found by Jean Horobin, this could have been, in part, a sample-size issue coupled with differences in Inner Farne colony size (nearly twice the size in 1966 – 1968 than in the present study) and environmental factors. Nevertheless, they did find birds of age two within the colony, and their arrival times of age-two birds and age-three birds being 31 and 17 days later, respectively, than birds of age eight or older were consistent with the differences of median times of capture of breeding birds age two (26 days later than older birds) and age three (16 days later) in the present study.

*SECTION 4*

**Clutch size in different age groups of known-age birds**
For birds age two years where nest contents were recorded, clutch/brood sizes are given in Table S3.

**Table S3:** Clutch/brood sizes for age two birds where contents recorded.

| Clutch/brood size | Frequency |
|---|---|
| 1 | 3 |
| 2 | 5* |
| 3 | 2 |

*one nest with two eggs near hatching; one nest with one egg and one young chick.

**Table S4:** Clutch/brood sizes for nests of nesting birds age four or less (Young), or five or more (Old)

| | **Frequencies of different clutch/brood sizes** | | | |
|---|---|---|---|---|
| **Clutch/brood:** | **1** | **2** | **3** | **4** |
| **Young** | 25 | 42 | 2 | 0 |
| **Old** | 48 | 116 | 10 | 2 |

Fisher’s Exact test on count data of Table S4: P = 0.45

**Table S5:** Clutch/brood sizes for nest contents of nesting birds age two, three or four.

| | **Frequencies of different clutch/brood sizes** | | |
|---|---|---|---|
| **Clutch/brood:** | **1** | **2** | **3** |
| **Age two** | 3 | 5 | 2 |
| **Age three** | 9 | 18 | 0 |
| **Age four** | 13 | 19 | 0 |

Fisher's Exact Test for Count Data of Table S5: P = 0.1

*SECTION 5*

**Table S6:** Age distribution of natal and non-natal nesting birds caught up to and including 2019.

| Age (years) | Natal | Non-natal | Total |
|---|---|---|---|
| **2** | 12 | 2 | 14 |
| **3** | 19 | 8 | 27 |
| **4** | 34 | 1 | 35 |
| **5** | 29 | 2 | 31 |
| **6** | 18 | 5 | 23 |
| **7** | 21 | 2 | 23 |
| **8** | 12 | 1 | 13 |
| **9** | 15 | 3 | 18 |
| **10** | 14 | 9 | 23 |
| **11** | 15 | 1 | 16 |
| **12** | 10 | 1 | 11 |
| **13** | 2 | 4 | 6 |
| **14** | 9 | 1 | 8 |
| **15** | 3 | 0 | 3 |
| **16** | 2 | 2 | 4 |
| **17** | 3 | 0 | 3 |
| **18** | 1 | 1 | 2 |
| **19** | 3 | 0 | 3 |
| **20** | 0 | 0 | 0 |
| **21** | 1 | 0 | 1 |
| **22** | 1 | 0 | 1 |

*SECTION 6*

**Figure S1:** Birds not caught on the nest: head-length distribution by age and natal status. Distributions of head lengths for known-age Arctic Terns caught 'randomly' (not caught on the nest) on Inner Farne or Brownsman. Each panel is a histogram of data with normal curves for head lengths constrained by means and standard deviations (sd) for female (red) or male (blue) Arctic Terns (females 70.2 mm, sd = 2.51; males 72.7 mm, sd = 1.68. (Fletcher & Hamer 2003)). Curve areas were estimated from the mixture proportions of females and males using the *normalmixEM* function from the R package *mixtools.* **a**, birds age four or less; **b**, birds age 5 or more; **c**, non-natal birds; **d**, birds caught on their natal site.

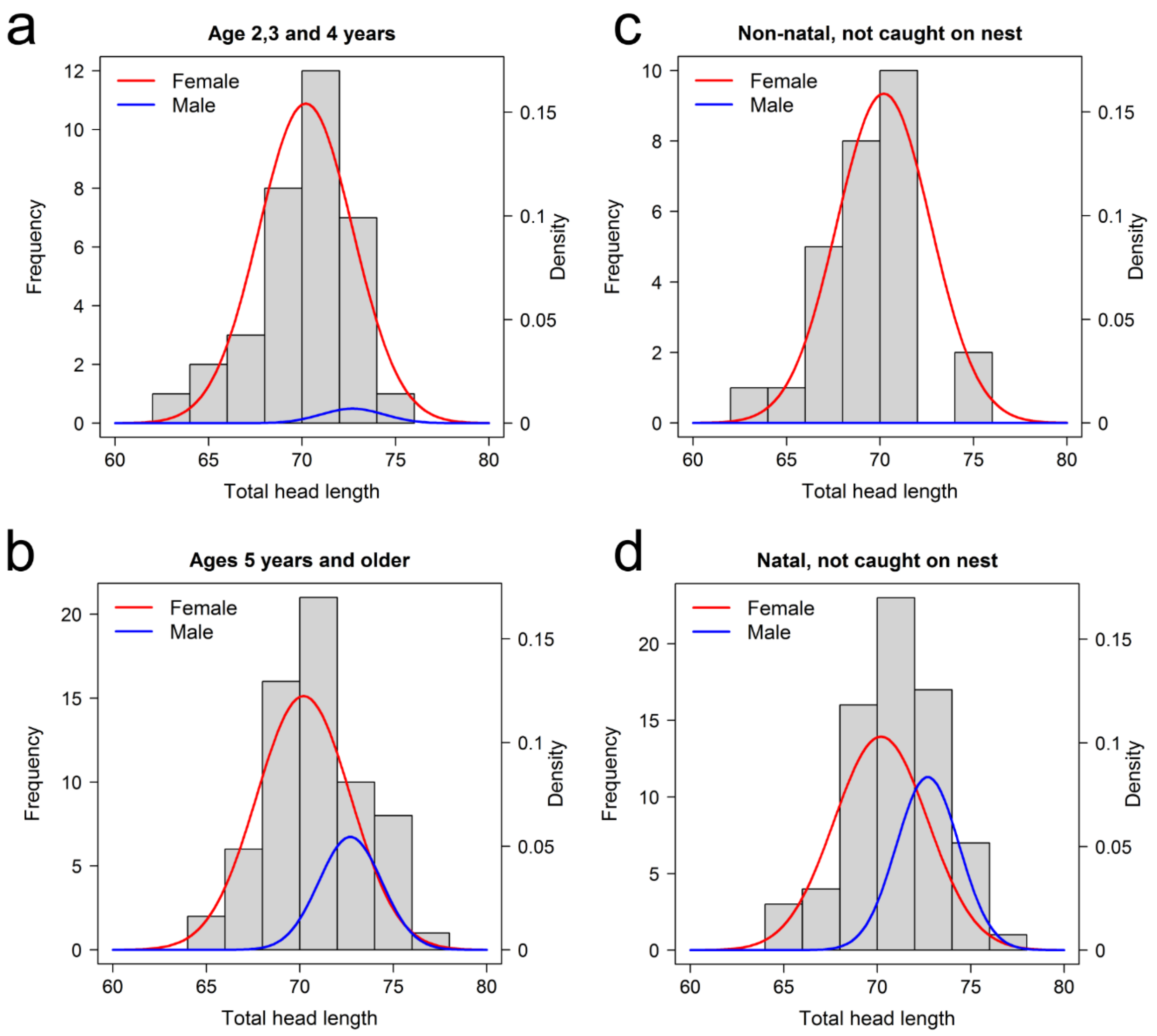

*SECTION 7*

**Table S7:** Coquet 2020-2022: age distribution of natal and non-natal birds

| Age | Natal | Non-natal |
|---|---|---|
| **3** | 0 | 3 |
| **4** | 0 | 2 |
| **5** | 3 | 0 |
| **6** | 0 | 0 |
| **7** | 1 | 4 |
| **8** | 1 | 3 |
| **9** | 0 | 3 |
| **10** | 0 | 1 |
| **11** | 0 | 0 |
| **12** | 1 | 0 |
| **13** | 0 | 0 |
| **14** | 1 | 0 |
| **15** | 0 | 0 |
| **16** | 0 | 0 |
| **17** | 0 | 0 |
| **18** | 0 | 0 |
| **19** | 1 | 2 |
| **20** | 0 | 0 |
| **21** | 0 | 0 |
| **22** | 1 | 0 |

*SECTION 8*

**Figure S2:** Coquet 2020-2022: head-length distributions by age class
Distributions of head lengths for nesting known-age Arctic Terns caught on Coquet Island in 2020 – 2022. Details as Fig. S2. **a**, birds age four or less; **b**, birds age 5 or more.

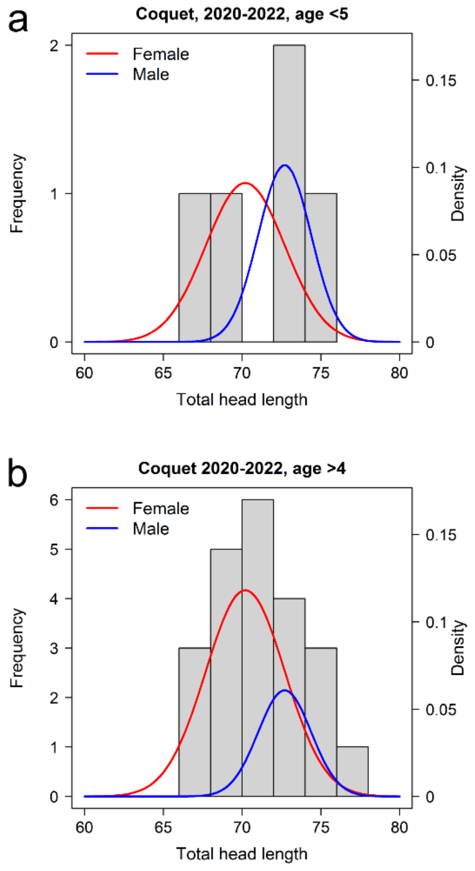
